# The Pleistocene species pump past its prime: evidence from European butterfly sister species

**DOI:** 10.1101/2020.09.04.282962

**Authors:** Sam Ebdon, Dominik R. Laetsch, Leonardo Dapporto, Alexander Hayward, Michael G. Ritchie, Vlad Dincă, Roger Vila, Konrad Lohse

## Abstract

The Pleistocene glacial cycles had a profound impact on the ranges and genetic make-up of organisms. Whilst it is clear that the contact zones that have been described for many sister taxa are secondary and have formed during the last interglacial, it is unclear when the taxa involved began to diverge. Previous estimates based on small numbers of loci are unreliable given the stochasticity of genetic drift and the contrasting effects of incomplete lineage sorting and gene flow on gene divergence. Here we use genome-wide transcriptome data to estimate divergence for 18 sister species pairs of European butterflies showing either sympatric or contact zone distributions. We find that in most cases species divergence predates the mid-Pleistocene transition or even the entire Pleistocene period. We also show that although post divergence gene flow is restricted to contact zone pairs, they are not systematically younger than sympatric pairs. This suggests that contact zones are not limited to the embryonic stages of the speciation process, but can involve notably old taxa. Finally, we show that mitochondrial and nuclear divergence are only weakly correlated and mitochondrial divergence is higher for contact-zone pairs. This suggests a possible role of selective sweeps affecting mitochondrial variation in maintaining contact zones.

**Impact Summary:** The influence of the Pleistocene glacial cycles on structuring species and genetic diversity in temperate taxa has permeated biogeographic and phylogeographic thinking for decades. Although phylogeographic studies have repeatedly claimed that the Pleistocene acted as a species pump, systematic tests of this idea based on robust estimates of species divergence are lacking. Here we estimate divergence times for all sister species pairs of European butterfly using genome-wide transcriptome data. We find that most species pairs are substantially older than the onset of Pleistocene glacial cycling. We also show that post divergence gene flow is restricted to pairs that form contact-zones. However, in contrast to expectations under a null model of allopatric speciation contract zone pairs are not necessarily younger than sympatric pairs.

## Introduction

Divergence in allopatry provides a simple null model of speciation (Mayr, 1947). Following geographic isolation and given enough time, reproductive isolation is inevitable as incompatibilities will eventually become fixed as a result of genetic drift and/or selection (Bateson, 1909, Dobzhansky, 1937, Muller, 1942). Taxa that evolved partial reproductive isolation in allopatry may come into secondary contact as a result of range shifts and – depending on their degree of reproductive isolation and niche overlap – either form a contact zone or invade each other’s range (Barton & Hewitt, 1985, Pigot & Tobias, 2013). If allopatric divergence dominates speciation, then local alpha diversity for a given clade cannot accrue until secondary sympatry is achieved (Weir & Price, 2011). Thus the forces that facilitate or hamper secondary sympatry and the timescale over which this occurs have profound consequences both for speciation and the spatial distribution of species diversity. While modern ranges only provide a snapshot of the dynamic history of range shifts, understanding the extent to which current range overlap between closely related species can be explained by their speciation history and *vice versa* has been at the core of speciation research (Coyne & Orr, 2004).

The glacial cycles of the Pleistocene had a profound effect on current diversity of temperate ecosystems (Hewitt, 1996, 2001, Hofreiter & Stewart). Populations of temperate taxa in Europe were isolated in ice-free refugia around the Mediterranean basin (Iberia, Italy, the Balkans and the larger Mediterranean islands) as glaciers encroached. The observation that the geographic ranges of many young taxa are restricted to individual glacial refugia in southern Europe (Dennis *et al.*, 1991, Hewitt, 1996, 1999, 2011) suggests that this repeated separation into and expansion out of glacial refugia has played a major role in their origin. The availability of allozyme and mitochondrial *(mt*) data in the 80s and 90s has spurred an abundance of case studies on intra- and interspecific diversity of European taxa including detailed investigations of hybrid zones in taxa ranging from fire-bellied toads (Kruuk *et al.*, 1999), the house mouse (Boursot *et al.*, 1996), grasshoppers (Barton, 1980, Butlin & Hewitt, 1985) to plants (Bacilieri et al., 1996) and marine mussels (Skibinski & Beardmore, 1979). The pervading evidence from these studies is that genetic diversity within and in, many cases, divergence between species is structured by refugia (Dapporto et al., 2019, Hewitt, 1996, Schmitt, 2007a).

### When was divergence between sister species initiated?

While it is clear that the hybrid zones we observe today are secondary contacts that formed after the last glacial maximum and may have formed many times over throughout the Pleistocene, it is far from clear when divergence between the sister taxa involved was initiated. One possibility is that the Pleistocene glacial cycles initiated species divergence directly by separating populations into allopatric refugia (i.e. a ‘species pump’ *sensu* Haffer (1969)). Another possibility is that the initial divergence between sister species predates the Pleistocene, and so, any build-up of reproductive isolation during the Pleistocene (e.g. via the fixation of intrinsic incompatibilities and/or reinforcement) occurred in populations that were already partially diverged. If the Pleistocene species pump hypothesis is correct, we would expect sister species divergence times to be concentrated during or at the beginning of the mid-Pleistocene transition 0.81.2 million years ago (MYA) which marks the onset of continent-wide glacial cycling (Bishop *et al.*, 2011). The idea that Pleistocene divergence acted as a species pump was first proposed in the context of American faunas (Avise *et al.*, 1998, Bernatchez & Wilson, 1998, Haffer, 1969), but has also dominated phylogeographic studies on European sister taxa (e.g. Habel *et al.*, 2008, Hewitt, 2000, 1996, Schmitt, 2007b, Schoville *et al.*, 2012). In contrast, other studies including some of the early work on European contact zones (Barton & Hewitt, 1985, Butlin & Hewitt, 1985) conclude that the taxa involved in such secondary contacts may substantially predate the Pleistocene (Abbott *et al.*, 2000, Hewitt, 1996, Klicka & Zink, 1997, Spooner & Ritchie, 2006). Thus it remains unclear to what extent divergence between sister taxa was initiated by ‘Pleistocene species pump’ dynamics or has an older, deeper origin?

A corollary for the hypothesis of allopatric speciation in different refugia is that range overlap is secondary. Since species can more easily invade each others ranges once sufficient premating barriers and ecological differentiation have developed, we would expect species pairs with overlapping ranges to be older overall than those without range overlap, all else being equal (Coyne & Orr, 2004). Support for this prediction comes from comparative studies showing that the proportion of range overlap (degree of sympatry (Chesser & Zink, 1994)) is positively (albeit weakly) correlated with genetic divergence (Barraclough & Vogler, 2000, Pigot & Tobias, 2013). However, a recent study in *Chorthippus* grasshoppers shows that subspecies that hybridise across contact zones can be older than currently sympatric species (Nolen *et al.*, 2020).

### Mito-nuclear discordance

Age estimates for recently diverged taxa have largely relied on single locus phylogenies and ignored incomplete lineage sorting. Hewitt (2011) summarises age estimates for European hybrid zones taxa including mammals, insects, amphibians, and reptiles, which range from hundreds of thousands to several million years ago. However, given that these estimates are based on different markers and calibrations, the extent to which glacial cycles have initiated speciation events remains unknown. Estimates based on mitochondrial *(mt*) data are particularly unreliable for at least three reasons. First, the mutation rate of mtDNA is highly erratic (Galtier *et al.*, 2009). Second, given the stochasticity of coalescence, the ancestry of a single locus (however well resolved) is a very poor measure of species divergence. In the absence of gene-flow divergence at a single locus may substantially predate the onset of species divergence, while it may be much more recent in the presence of gene flow (Knowles & Carstens, 2007, Wang & Hey, 2010). Mito-nuclear discordance in both directions has been found in a large number of animal systems (Toews & Brelsford, 2012) including several closely related species of European butterflies (Dincă *et al.*, 2019, Hinojosa *et al.*, 2019, Wiemers *et al.*, 2010). Finally, mtDNA does not evolve neutrally since transmission of mitochondria is completely linked to maternal inheritance of endosymbionts such as *Wolbachia* and *Spiroplasma* and, in organisms with Z/W sex determination, of the W chromosome. Thus *mt* diversity and divergence may be driven largely by selective sweeps (including introgression sweeps) rather than neutral gene flow and genetic drift (Galtier *et al.*, 2009, Hurst & Jiggins, 2005, Jiggins, 2003, Martin *et al.*, 2020).

### European butterflies as a model group

Testing whether climate-induced Pleistocene range shifts have triggered speciation or patterned older splits between species requires replication both at the level of genetic loci and at the level of speciation events. Although we can now generate WGS data for any species, there are surprisingly few reliable estimates for the onset of divergence between European sister species and such estimates are lacking even for well-studied contact zone taxa (but see Nolen *et al.*, 2020, Nürnberger *et al.*, 2016).

Lepidoptera are arguably the best-studied arthropod family: European butterflies provide a unique opportunity to investigate divergence and speciation processes comparatively (Dapporto *et al.*, 2019). Near-complete information on geographic ranges and key life-history traits (e.g. voltinism and host plant range) is available (Kudrna, 2019, Tolman & Lewington, 2013). Additionally, the taxonomy of all 496 European species (Wiemers *et al.*, 2018) is well resolved and a complete, multilocus phylogeny of all European taxa exists (Dapporto *et al.*, 2019). This combined with extensive DNA barcode reference libraries (Dapporto *et al.*, 2019, Dincă *et al.*, 2015), facilitates the identification of species (especially in the case of cryptic taxa) and provides extensive sampling of sister species pairs, many of which abut at narrow contact zones (Dennis *et al.*, 1991, Platania *et al.*, 2020, Vodă *et al.*, 2015) (Figure 1). Secondary contact zones have been described in detail for several European taxa, including *Spialia orbifer* and S. *sertorius* (Lorkovic, 1973), the Italian *Pontia* hybrid zone (Porter *et al.*, 1997) and the contacts between *Iphiclides podalirus* and *I. feisthamelii* and between *Melanargia galathea* and *M. lachesis* along the Pyrenees (Gaunet *et al.*, 2019b, Habel *et al.*, 2017, Wohlfahrt, 1996).

**Figure 1:**
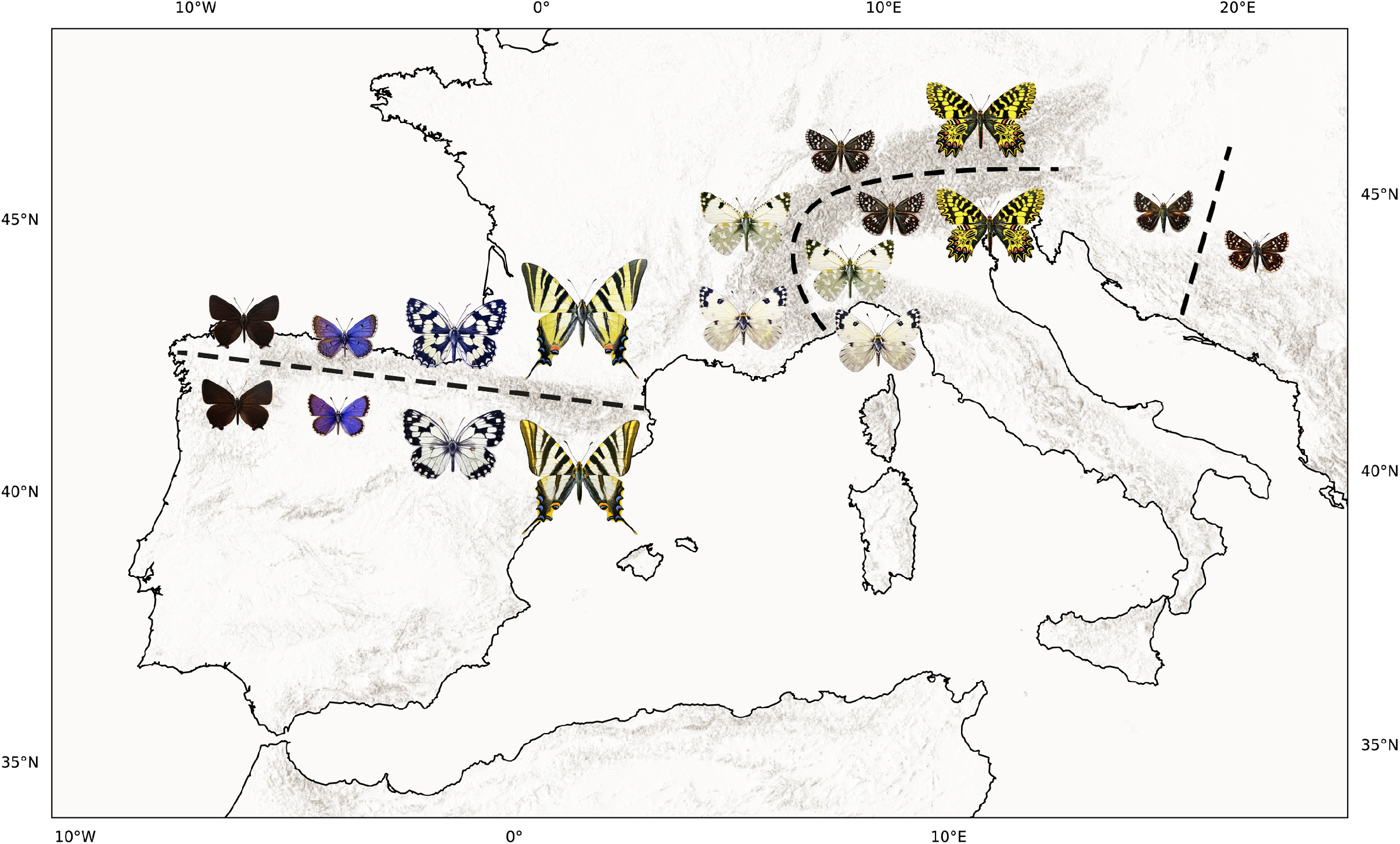
Nine of the 18 sister species pairs of butterfly in which we quantified genome-wide divergence meet at contact zones in southern Europe. In the left group, from left to right across northern Iberia are *Satyrium, Pseudophilot.es, Melanargia,* and *Iphiclides.* In the center group, from bottom to top across the Alps are *Pontia, Euchloe, Pyrgus,* and *Zerynthia.* Finally, on the right across the Balkans is the genus *Spialia.*

Here we use European butterflies as a model system to investigate to what extent the divergence times between sister species in this group are concentrated in the Pleistocene, as predicted by the Pleistocene species pump hypothesis, and test how well recent sister species fit a null model of divergence in allopatry. Although European butterflies have been studied intensively, with few exceptions (see Talla *et al.*, 2017) the robust estimates of divergence required for any systematic comparison of speciation are lacking. We generate RNAseq data for 18 sister species pairs and ask the following specific questions:

i. Has speciation been initiated during the Pleistocene as envisaged by the species-pump hypothesis or did the glacial cycles pattern pre-existing, older subdivisions?
ii. Are sister species pairs that form contact zones younger than pairs that overlap in range?
iii. Is there evidence for gene flow in contact zone species?
iv. How strongly correlated are mitochondrial and nuclear divergence and do contact zone pairs show increased mito-nuclear discordance?

## Methods

### Sampling and molecular work

We identified true sister species pairs in the European butterfly phylogeny (Dapporto *et al.*, 2019). Species pairs involving island and mountain endemics were excluded, as these cannot achieve secondary sympatry. We also did not consider species pairs that are unlikely to have originated in Europe, e.g. sister pairs involving North American taxa. Following these criteria, we sampled 18 sister species pairs. Our sampling includes 7.3 % of European butterfly species and almost all ‘good’ butterfly sister species pairs in Europe (Descimon & Mallet, 2009).

Field sampling was conducted over multiple seasons (2016-2019) at several locations across Southern and Central Europe (Portugal, Spain, France, Hungary, Romania). Samples were hand-netted in the field, flash-frozen in a liquid nitrogen dry shipper (Voyageur 12) and stored at −70 °*C* shortly after capture (wings were retained for identification). Specimen identifications were confirmed for 22 samples by DNA barcoding using LepF/R primers (Hajibabaei *et al.*, 2006) and existing reference databases (Dincă *et al.*, 2015). We were unable to obtain fresh material for *Erebia euryale* and *E. ligea,* and *Fabriciana adippe* and *F. niobe* (two remaining sister pairs meeting our sampling criteria).

RNA extractions were prepared by dividing individuals bilaterally and using one side. RNA was extracted following a hybrid protocol by homogenising samples with TRIzol, then digesting DNA and eluting RNA using the Purelink RNA Purification kit protocol. Extracted RNA was submitted to Edinburgh Genomics to generate automated TruSeq stranded mRNA-seq libraries. Libraries were sequenced on an Illumina NovaSeq platform using 100PE reads after polyA selection. For each species, where possible, we generated RNASeq data for two samples, one male and one female from different localities. Transcriptome data for 66 samples (across 38 species) were generated and analysed previously by Mackintosh *et al.* (2019). Of these, 26 samples from 13 species are included in the present analysis (Table S1).

### Generating transcriptome assemblies

Reads were processed following the pipeline developed by Mackintosh *et al.* (2019). Reads were trimmed and checked for quality using FastQC v0.11.8 (Andrews *et al.*, 2010) both before and after trimming with FastP v0.20.0 (Chen *et al.*, 2018) using MultiQC v1.7 (Ewels *et al.*, 2016) to visualise the results. Trimmed reads were assembled into *de novo* transcriptomes using Trinity v2.8.5 (Grabherr *et al.*, 2011), pooling data-sets by species.

Transcriptome completeness was assessed using BUSCO v3 (Simão *et al.*, 2015) with the *insectaodb9* database. Transcripts were processed with Transdecoder v5.5 (Haas *et al.*, 2016), and retained based on BLAST (Camacho *et al.*, 2009) and HMMER (Finn *et al.*, 2011) homology search results. Read pairs from each sample were mapped against respective species transcriptome, composed of the longest isoform of each complete protein-coding transcript, using BWA MEM (Li, 2013). Coverage at mapped sites was determined using GATK CallableLoci v3.5 (McKenna *et al.*, 2010). Sites with at least 10 fold coverage and a minimum mapping quality of 1 in each sample were considered suitable for variant calling. Callable loci were intersected between individuals using BEDTools v2.28 (Quinlan & Hall, 2010), variants were called using FreeBayes v1.3.1 (Garrison & Marth, 2012) and filtered for unbalanced SNPs and missing genotypes (RPL ≥1 RPR≥1 SAF≥1 SAR?1 N_MISSING=0) using BCFtools filter v0.1.19 (Li, 2011).

To generate comparable data-sets across all samples, Orthofinder v2.3.3 (Emms & Kelly, 2015) was used to cluster proteins into orthogroups. Orthogroups were labelled single-copy orthologues (SCOs) if one protein of each taxon was present. Genus single-copy orthologues (GSCOs) were diagnosed based on the presence of single-copy proteins within the focal pair. Protein sequences from each orthogroup were used to align equivalent DNA sequences using Translatorx v12.0 (Abascal *et al.*, 2010).

Data were generated for 36 species (18 sister pairs) from five families. For 17 pairs, data were generated from 665 SCOs from high-quality transcriptomes (BUSCO scores > 90%). For the pair of *Zerynthia* species (one of which, *Zerynthia polyxena,* was sampled as a larva) GSCOs (5000 orthologues) were used to avoid restricting the SCOs for other pairs. With the exception of the *Zerynthia* pair, all analyses are based on SCO to enforce consistent comparisons across pairs. While the SCO data-set is much smaller than the pair GSCO data-sets and likely enriched for conserved and highly expressed genes, this has very little impact on estimates of divergence and diversity at fourfold degenerate (4D) sites, as these are highly correlated (Figure S1 and Mackintosh *et al.* (2019)).

### Estimating gene and population divergence

For each species pair, we calculated *d_xy_* at 4D sites using sequence alignments for one or two diploid samples from each species. This calculation is implemented in the script orthodiver.py (www.github.com/samebdon/orthodiver).

Information on generation times was compiled from Collins Butterfly Guide (Tolman & Lewington, 2013) (Table 1). For species in which generation times vary with latitude, we assumed the minimum generation time of the southern part of the range. This is a reasonable long term average, given that European glacial refugia are located around the Mediterranean.

**Table 1:**
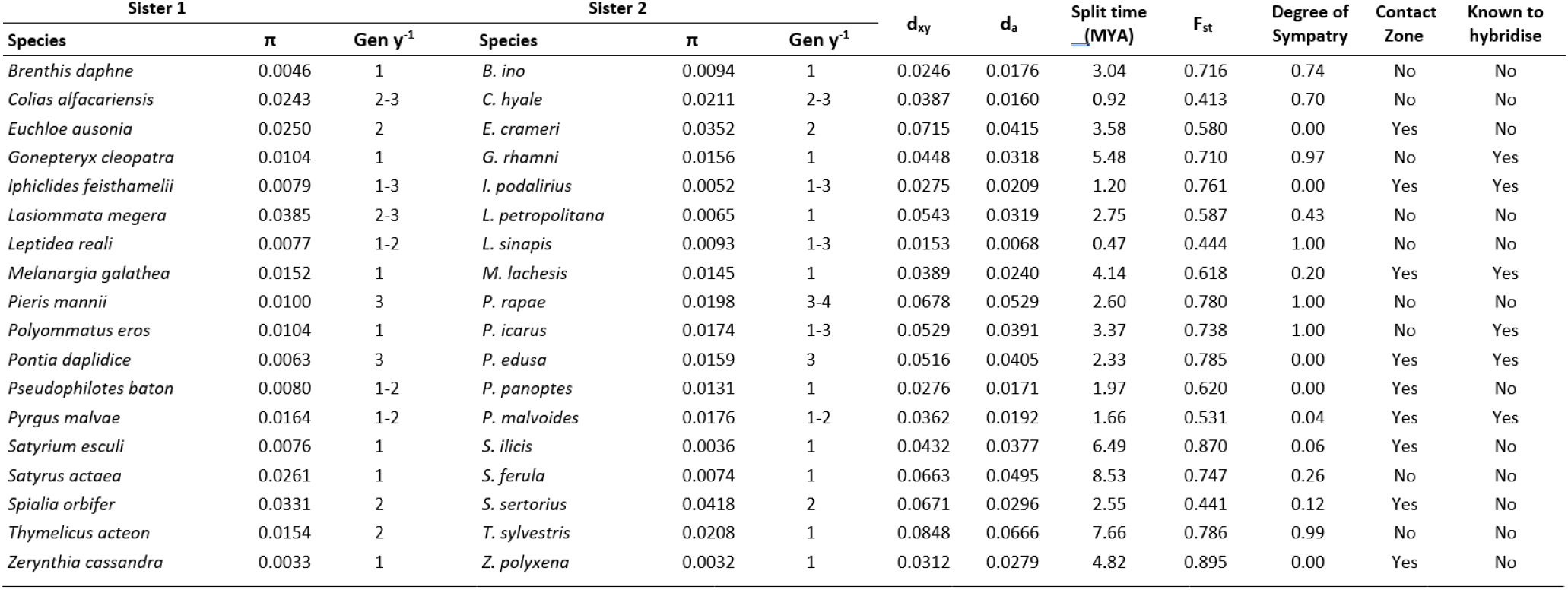
Estimates of mean gene divergence (*d_xy_*), net gene divergence (*d_a_*) and differentiation (*F_st_*) at 4D sites and lower bounds for species divergence times for 18 sister species pairs of European butterfly. Gen *y*^−1^ is the number of generations per year.

We considered the distribution of pairwise differences in blocks of a fixed length of 4D sites. The block size for each pair was chosen to give an average of three pairwise differences between sister species per block. To test whether species pairs show evidence for gene flow, we compared the observed distributions to analytic expectations under a model of strict divergence without gene flow (given estimates of *T* and ancestral *N_e_* obtained from *d_a_* and mean *π*).

In the absence of recombination within blocks, analytic expectations for the distribution of pairwise differences are available (Lohse *et al.*, 2011, Wilkinson-Herbots, 2012). However, given the high rate of recombination (relative to mutation) in butterflies (Martin *et al.*, 2019, 2016) and the substantial span of 4D blocks, we expect the empirical distribution to be narrower than this analytic expectation. We re-sampled (without replacement) 10,000 data-sets of equal size as the observed data-sets from the expected distribution of each species and tested whether the likelihood of the observed distribution of pairwise differences falls within the distribution of likelihoods obtained from re-sampled data-sets.

### Estimating range size and overlap

Geographic ranges were quantified as follows: we obtained occurrence data over Europe for all focal species with a resolution of 60’ latitude and 30’ longitude by critically revising the data from the Distribution Atlas of European Butterflies and Skippers (Kudrna *et al.*, 2011) and by adding data from Roger Vila’s collection stored at Institut de Biologia Evolutiva (Barcelona). To calculate range overlap we applied the biodecrypt function (Platania *et al.*, 2020) of the recluster R package (Dapporto *et al.*, 2013). This function computes alpha hull with a given concavity (*α*) and evaluates the area of overlap among pairs of species. We used *α* = 2 and *α* = 3 for species with discontinuous and continuous distributions in Europe respectively. We quantified the range overlap of each species pair and calculated the degree of sympatry as:

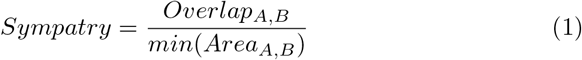
that is the fraction of the smaller range involved in the overlap. In the following, we consider sister pairs with a degree of sympatry ≤ 0.2 contact zone pairs and those with a degree of sympatry > 0.2 sympatric. Based on this, we classified nine pairs as contact zone taxa. However, since there are only two species pairs with intermediate levels of sympatry (> 0.2 and < 0.7), our comparisons of contact zone and sympatric pairs are robust to a wide range of thresholds.

### Mitochondrial diversity and divergence

Sequence alignments for the COI barcode locus were obtained from the BOLD database (Ratnasingham & Hebert, 2007) for all 18 sister species pairs. Sequence alignments are deposited in the dryad repository xxx. For each species, we included all available sequence records from Europe. Mean pairwise diversity (π) within species and divergence (*d_xy_*) across all sites were computed using DnaSP (Librado & Rozas, 2009).

We obtained the average gene divergence time for each pair from the multilocus calibrated phylogeny of European butterflies of Wiemers *et al.* (2020) as half of patristic distances calculated with distTips function of the adephylo R package (Jombart & Dray, 2010). The correlation between our estimates of species divergences and these node ages was explored with standardized major axis (SMA) regression, using the ‘sma’ function of the ‘smatr’ R package. SMA estimates slope and intercept and tests if slope differs from one.

## Results

### Most European butterfly sister species predate the Pleistocene

We assumed a simple null model of divergence without gene flow, neutrality and an infinite sites mutation model and used net mean divergence at fourfold degenerate (4D) sites *d_a_* = *d_xy_* – π (Nei & Li, 1979) to estimate species divergence time 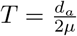. Here *μ* is the *de novo* mutation rate per generation (per base). We assumed *μ* = 2.9 * 10^−9^, an estimate of the spontaneous mutation rate obtained from parent-offspring trios of South American *Heliconius melpomene* butterflies (Keightley *et al.*, 2014b). Since both violations of the mutation model (back-mutations) and the demographic model (gene flow) reduce *d_a_*, this time estimate is a lower bound of the true species divergence time. We converted estimates of species divergence time (*T*) into years (*τ*) using the mean generation time of each pair (Table 1).

Species divergence times obtained from *d_a_* at fourfold degenerate sites (4D) ranged from 0.47 (*Leptidea*) to 8.5 (*Satyrus*) MYA, with a mean of 3.8 MYA, (Figure 2). Even though these point estimates are lower bounds of species divergence (see Discussion), they not only substantially predate the mid-Pleistocene transition (15 out of 18 pairs) but, in the majority of cases (11 out of 18 pairs), are older than the entire Quaternary period ≈ 2.6 MY (Table 1). Of the seven taxa with Pleistocene *τ* estimates, three fall in the early Pleistocene: *Pseudophilotes* (1.97 MYA), *Pontia* (2.33 MYA) and *Spialia* (2.55 MYA).

**Figure 2:**
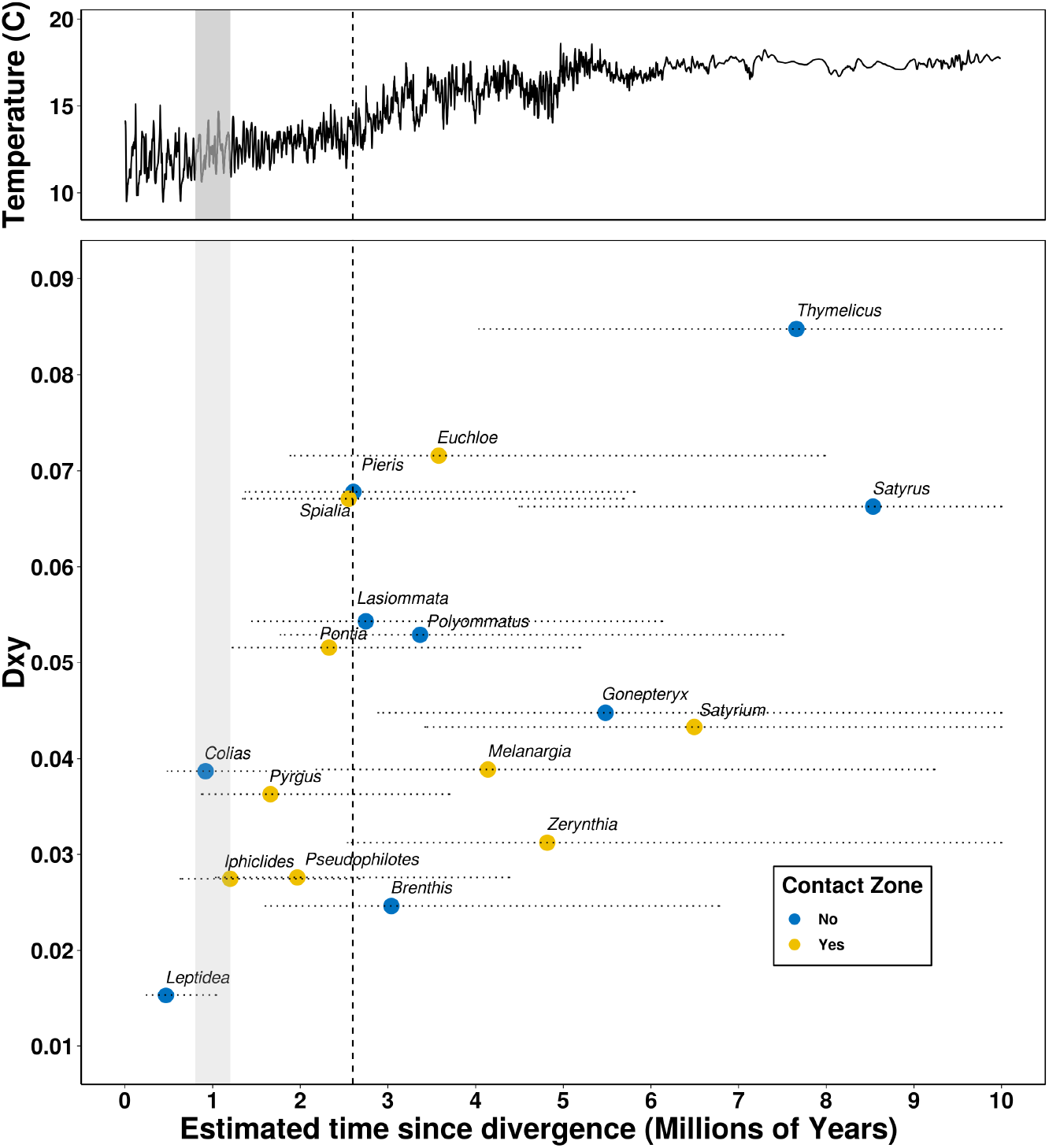
Species divergence time estimates (*τ*) plotted against mean genetic divergence (*d_xy_*) for 18 European butterfly sister species pairs. Pairs which abut at contact zones (degree of sympatry ≤ 0.2) are shown in yellow, sympatric pairs with substantial range overlap (> 0.2) in blue. The vertical dashed line represents the beginning of the Pleistocene (2.6 MYA), the vertical grey bar indicates the mid-Pleistocene transition (0.8 to 1.2 MYA). The horizontal dotted lines represent the 95% confidence intervals of *τ* estimates given the uncertainty in the mutation rate used for calibration (1.3-5.9×10^−9^ (Keightley *et al.*, 2014b)). The temperature data (five-point running means of global surface temperature) are taken from Hansen *et al.* (2013)

Even accounting for the considerable uncertainty in the mutation rate estimate we used in our calibration (Keightley *et al.*, 2014b), only five species pairs show 95 % CI of τ that overlap the mid-Pleistocene transition.

### Sister pairs that form contact zones are not significantly younger than sympatric pairs

Mean gene divergence (*d_xy_*) at 4D sites between sister species ranged from 1.5% to 8.5%, with a mean of 4.7% (Table 1, Figure 2) across the 18 pairs. There are two reasons to expect species pairs that form contact zones to be younger than sympatric pairs: First, if speciation under a null model of divergence in allopatry is initiated by periods of vicariance, the formation of a contact zone (parapatry) represents an earlier stage in the transition to complete reproductive isolation and substantial range overlap (sympatry) (Coyne & Orr, 2004). Second, any gene flow across contact zones would reduce *d_a_* and hence our estimate of species divergence. The nine pairs that form contact zones (degree of sympatry ≤ 0.2) have a lower net divergence (*d_a_* = 0.0287, SD = 0.00930) than the nine sympatric pairs (degree of sympatry > 0.2, *d_a_* = 0.0347, SD = 0.0195 Table 1), however, this difference is not significant (t = −0.82999, df = 11.478, p = 0.210). Additionally, we find no relationship between the degree of sympatry and *d_a_* (t = 0.723, df = 16, p = 0.480). Similarly, we may expect pairs that are still able to form hybrids (i.e. for which F1s have been observed in the wild) to be younger than those that do not. However, contrary to this expectation, we again find no significant difference in net divergence between pairs which do and do not hybridise (*d_a_* 0.0293 and 0.0329 respectively, t = −0.582, df = 15.861, p = 0.284).

### Evidence for recent gene flow in some contact zone pairs

Rather than fit explicit models of demographic history which is difficult using transcriptome data for minimal samples of individuals, we tested for signals of post divergence gene flow in the distribution of pairwise differences in sequence blocks of a fixed length. This distribution may differ from analytic expectations under a model of neutral divergence (and assuming no recombination within blocks (Lohse *et al.*, 2011, Wilkinson-Herbots, 2012)) in two ways: while gene flow widens the distribution of pairwise differences, recombination within blocks narrows it (Wall, 2003). Thus, in the absence of gene flow, we would expect empirical distributions to be narrower than the analytic expectation while wider distributions are indicative of post-divergence gene flow.

The empirical distribution of pairwise differences deviated significantly from the expectation in a majority of species pairs (12 out of 18) (Figure 3 & S6). Of these, eight pairs have narrower distributions than expected, compatible with recombination within blocks and four pairs have wider distributions than expected, compatible with post-divergence gene flow (*Pseudophilotes, Pontia, Iphiclides, Zerynthia*). While the eight pairs with narrower distributions are equally split between contact and sympatric pairs, all four taxa with wider distributions are contact zone pairs (Figure 3). However, given the limited number of pairs overall, this difference between contact zones and sympatric pairs is not significant (Fisher’s exact test, p = 0.0901).

**Figure 3:**
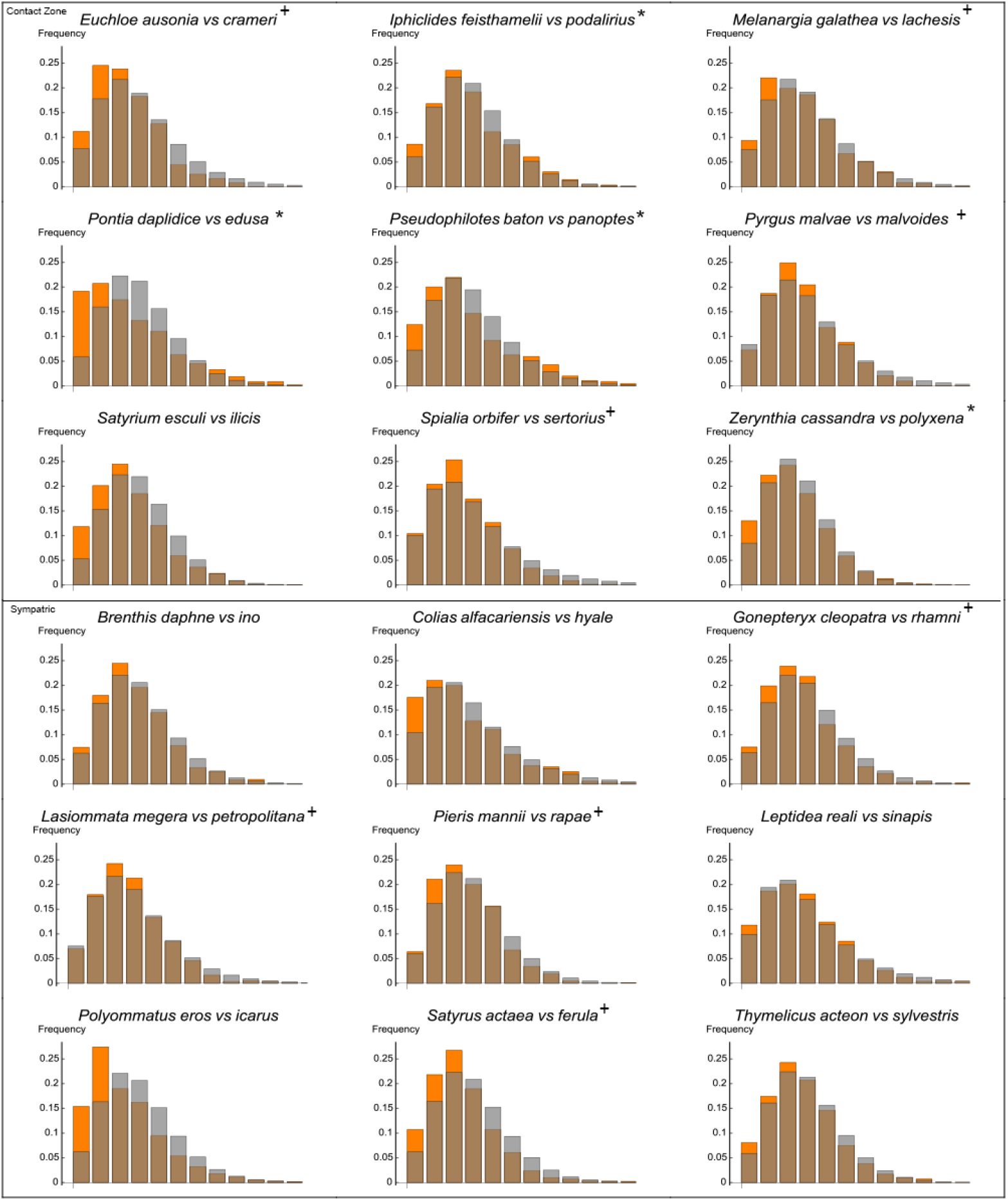
The distribution of pairwise differences (in blocks of a fixed length of 4D sites) in contact (upper box) and sympatric (lower box) pairs. The observed distribution in single-copy orthologues is shown in orange, the expectation under a history of strict divergence (estimated from π and *d_a_*) in grey. Pairs that show wider than expected distributions are marked with an asterisk (*) and species which show narrower than expected distributions are marked with a plus (+).

### Pervasive mito-nuclear discordance in contact zone species pairs

Our estimates of species divergence are based on average net divergence (*d_a_*) across many hundreds of genes and are robust to how orthologues are filtered (Figure S1). Given that previous studies on European butterflies have been largely based on mitochondrial (*mt*) phylogenies, an important question is to what extent *mt* divergence is correlated with mean nuclear divergence. We find that both *d_a_* and *d_xy_* at COI are positively but only weakly correlated with mean nuclear divergence (Figure 4). The correlation is weaker for *d_a_* than *d_xy_* (*R*^2^ = 0.27 and 0.31 respectively) which is compatible with mitochondrial diversity (and hence da) being disproportionately affected by selective sweeps. Similarly, comparing the relation between *mt* and nuclear da between contact zone and sympatric pairs, we find a much shallower slope for contact zone pairs (0.29 compared to 0.99, Figure 4)). This difference is largely a result of reduced *mt* diversity in contact zone compared to sympatric pairs (mean π = 0.0030, SD = 0.0014 and π = 0.0047, SD = 0.0031 respectively t = 1.5763, df = 11.324, p = 0.0712). This suggests that *mt* diversity is more strongly affected by selective sweeps in contact zone species than in sympatric pairs. We find no corresponding difference in nuclear diversity between contact zones and sympatric pairs (t = −0.0139, df = 31.539, p = 0.506) and, in general, no correlation between nuclear and *mt* diversity (Figure S3 and (Mackintosh *et al.*, 2019)).

**Figure 4:**
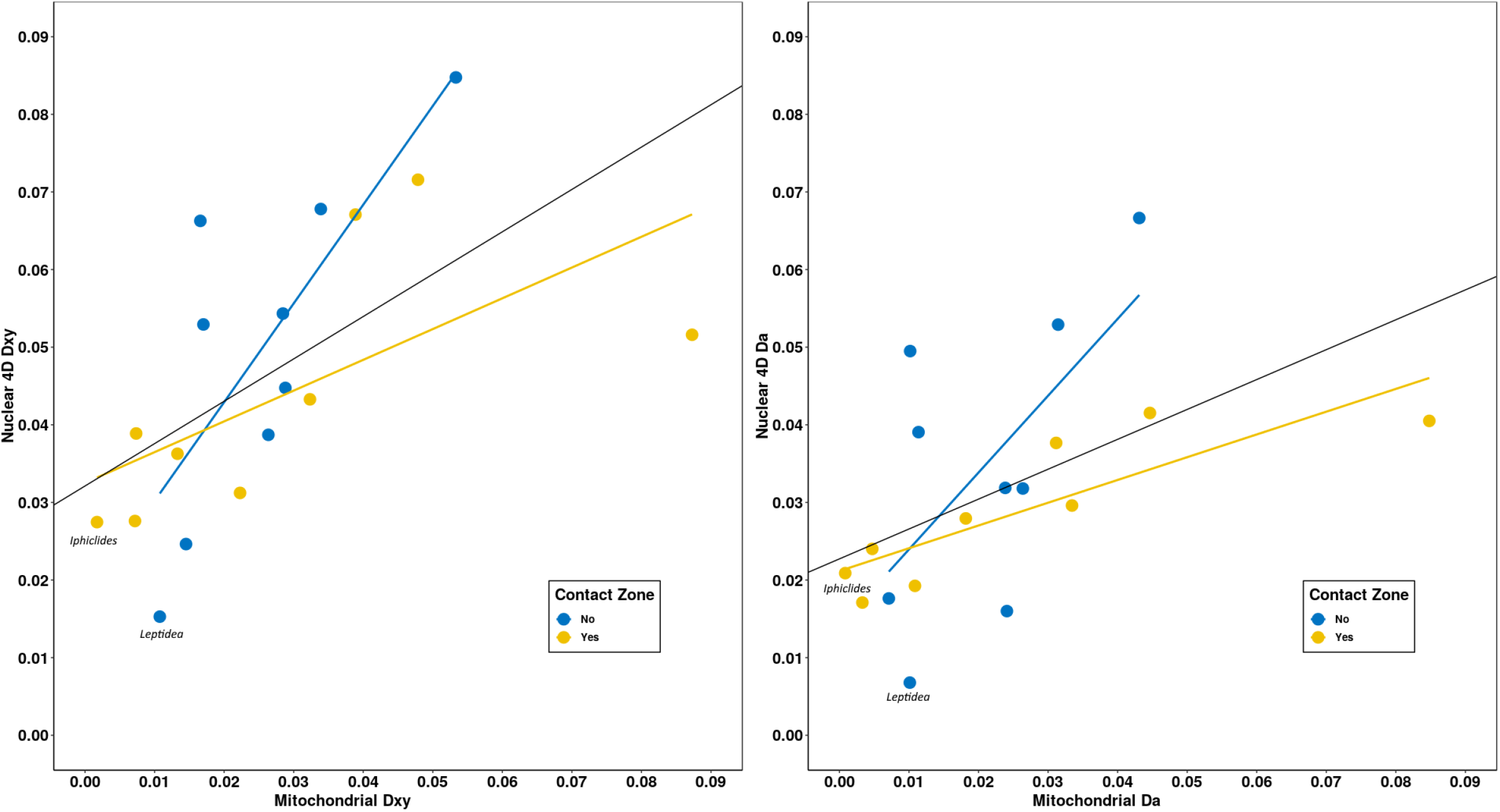
Mitochondrial *d_xy_* (left) and *d_a_* (right) are weakly correlated with mean nuclear divergence (*R*^2^ = 0.3565 and 0.2732, respectively). The coloured lines show the interactions for pairs that form contact zones and sympatric pairs. The two highlighted pairs (*Iphiclides* and *Leptidea)* have known *Wolbachia* associated sweeps and low *mt π* (and so high *mt d_a_*).

Our estimates for the lower bound of sister species divergence differ substantially from the ages of the corresponding nodes in the Wiemers *et al.* (2020) phylogeny for individual pairs (Fig S4). This is unsurprising given that the latter are largely informed by mtDNA data (Fig S5). However, perhaps surprisingly (given the difference in calibration, data and inference approach) our estimates are not consistently older or younger than the node ages of Wiemers *et al.* (2020) (*t_paired_* = −1.105, df = 17, p = 0.285). A standardized major axis regression shows a significant relationship (R squared = 0.3657, p = 0.00780), a slope (1.377) not different from one (r= 0.3786, p = 0.121) and an intercept (−0.5750) not different from zero (Fig S4).

### Genetic diversity does not correlate with relative range size

Genetic diversity at 4D sites within all 36 species ranged from 0.32% to 4.2% with a mean of 1.5%. Given the *H. melpomene* mutation rate of μ = 2.9×10^−9^ (Keightley *et al.*, 2014b), these correspond to effective population sizes ranging from 280,000 to 3,600,000 with a mean of 1,300,000. Mackintosh *et al.* (2019) tested whether neutral genetic diversity across European butterflies correlates with geographic range and found no significant relation across 38 taxa. Our sampling of species pairs allows for a simpler, alternative test of the potential relationship between diversity and range size using sister-clade comparisons which are less sensitive to potential phylogenetic correlates and uncertainty in current range estimates. If diversity is a function of range size, we expect the species in a pair with the larger range to have higher genetic diversity than the species with the smaller range. We indeed find a difference in the expected direction, 0.0167 (SD = 0.0114) vs. 0.0139 (SD = 0.00865), although the effect of relative range size is not significant (*t_paired_* = 1.127, df = 17, p = 0.138, Figure S2).

## Discussion

We quantify and compare genome-wide divergence across 18 sister species pairs of European butterfly. Simple lower bounds for the onset of species divergence based on net gene divergence (*d_a_*) and a direct mutation rate estimate for butterflies suggest that the majority of pairs have diverged before the onset of the major Pleistocene glacial cycles. Our results support the notion that the modern contact zones are secondary between species that began to diverge much earlier, in the Pliocene or early Pleistocene. Thus, even though the current ranges of many taxon pairs reflect glacial refugia, their initial divergence during the Pliocene or early Pleistocene is unlikely to have been triggered by repeated cycles of range connectivity and vicariance into refugia, as envisaged by the species-pump hypothesis and earlier phylogeographic studies based on *mt* and allozyme data (e.g. Habel *et al.*, 2008, Lai & Pullin, 2004, Schmitt, 2007b, Todisco *et al.*, 2010, Zinetti *et al.*, 2013) because substantial glaciation across continents did not develop (Bishop *et al.*, 2011) until the ‘mid-Pleistocene transition’ 0.8-1.2 MYA and the shift from 41,000 to 100,000 year glacial cycles. Given the antiquity of most sister species, it is unsurprising that we do not find any relationship between current range overlap and the time since divergence. Specifically, species pairs which form contact zones are not significantly younger than pairs that broadly overlap in range. However, we do find that strong signals of post-divergence gene flow are restricted to contact-zone pairs. It is likely that the absence of sympatric pairs with significant gene flow reflects a simple survivorship bias: any such pairs with significant gene flow might have already collapsed. Similarly, we are more likely to observe old contact zones pairs that have survived repeated glacial cycles.

Our finding that *mt* divergence between sister species is only weakly correlated with mean nuclear divergence and that net *mt* divergence is greater in contact zone than sympatric species pairs (as a result of reduced genetic diversity), suggests that the former are subject to more frequent selective sweeps linked to mitochondria. Such sweeps may be acting on *mt* variation directly or, indirectly, through maternally inherited genomes or chromosomes (e.g. *Wolbachia* (Jiggins, 2003) and the W chromosome) and have been documented in a number of Lepidopteran systems (Gaunet *et al.*, 2019a, Graham & Wilson, 2012, Kodandaramaiah *et al.*, 2013, Martin *et al.*, 2020, Ritter *et al.*, 2013). Our results raise the intriguing possibility that such sweeps could play a role in the build-up of reproductive isolation (Giordano *et al.*, 1997, Rokas, 2000, Shoemaker *et al.*, 1999).

### Sources of dating uncertainty and bias

Since we have assumed a simple demographic null model of species divergence without gene flow, our estimates of divergence between sister species should be interpreted as lower bounds. Any gene flow between sister species would reduce *d_a_* and species divergence estimates both by decreasing *d_xy_* and by potentially increasing π (in the recipient species).

Calibrating absolute split times involves assumptions about both the generation time and the mutation rate. We have assumed that the mutation rate is the same (per generation) across all species pairs, irrespective of their generation time and applied a direct lab estimate of the per generation mutation rate from the tropical butterfly *H. melpomene*. Whilst there is good evidence for a generation time effect on mutation rates in invertebrates (Thomas *et al.*, 2010), our assumption of a simple linear relationship between generation time and sequence divergence may be overly simplistic. In particular, if temperate European species, which have longer average generation times than *H. melpomene*, have a higher per generation mutation rate, we would have overestimated the age of sister species. In contrast, given that generation time varies between populations, species, and likely through time, our use of the average minimum generation time (within each pair) as a proxy for the long term generation time is conservative: assuming longer average generation times would yield even older estimates species divergence. Given these uncertainties in calibration, our absolute time estimates should be interpreted with caution until direct mutation rate estimates for temperate butterflies are available. However, in the absence of information on mutation rate heterogeneity across Lepidoptera, our main conclusion that divergence between most sister species of European butterflies predates the Pleistocene would still hold if mutation rates were higher by a factor of two. Given that the direct estimate of the *de novo* mutation rate in *H. melpomene* is similar to spontaneous mutation rate estimates for other insects (Keightley *et al.*, 2014a), this seems extremely unlikely. While our split time estimates may be surprising in light of previous phylogeographic studies on European butterflies based on *mt* diversity (e.g. Habel *et al.*, 2008, Lai & Pullin, 2004, Schmitt, 2007b, Todisco *et al.*, 2010, Zinetti *et al.*, 2013), our divergence estimate for *Leptidea reali* and *L. sinapis*, the youngest and only pair for which divergence has been estimated based on genome-wide data before, is lower than previous estimates (Talla *et al.*, 2017).

### Glacial cycling and the Messinian salinity crisis

Taking our estimates of species splits at face value, the species divergence for 10 species pairs predates the onset of Pleistocene glacial cycling > 2.6 MYA (Gibbard & Head, 2009). This is not compatible with the idea that, overall, speciation processes in European butterflies were initiated by the range shifts into and out of glacial refugia during the Pleistocene. However, our age estimates do of course not rule out that Pleistocene range shifts and vicariance may have played a role in completing speciation processes, e.g. through reinforcement and/or the evolution of intrinsic incompatibilities.

An event which may have contributed to speciation in Europe before the onset of Pleistocene glacial cycling is the Messinian salinity crisis (MSC) ≈ 6MYA during which the Mediterranean greatly reduced in size (Krijgsman *et al.*, 1999). As a consequence, Europe and Africa were connected across the strait of Gibraltar until the Zanclean flood when the Atlantic reconnected to the reexpanding Mediterranean sea. This must have created a strong dispersal barrier for many species that previously had continuous distributions around the Mediterranean basin and may have initiated the divergence into the east and west European/Mediterranean sister taxa. While the MSC has been considered as a plausible trigger of species divergence in amphibians (Nürnberger *et al.*, 2016) and reptiles (Kaliontzopoulou *et al.*, 2011), it has rarely been invoked for Lepidopterans (see recent insights into mitochondrial lineages in *Melitaea didyma* (Dincă *et al.*, 2019)) which have assumed to have been younger.

### Do European butterfly species fall within the grey zone of speciation?

Roux *et al.* (2016) conducted a comparative analysis of divergence and gene flow across 61 pairs of sister taxa and found that pairs with net synonymous divergence of > 2% rarely show evidence for ongoing gene flow. In contrast, taxa with *d_a_* between 0.5% and 2% may show some evidence for ongoing gene flow and ambiguous species status, suggesting that speciation may be incomplete. While our five youngest pairs (*Brenthis*, *Colias*, *Leptidea*, *Pseudophilotes*, and *Pyrgus*) fall in this “grey zone of speciation”, we only find evidence for gene flow in one (*Pseudophilotes*). In contrast, we find a clear gene flow signal in three more diverged pairs: *Iphiclides*, *d_a_* = 2.09%; *Zerynthia*, *d_a_* = 2.79%; *Pontia*, *d_a_* = 4.05%. However, as we have focused sampling on “good species” *sensu* Mallet (Descimon & Mallet, 2009) we are missing the recent (intraspecific) end of the continuum of divergence described by Roux *et al.* (2016). It will be interesting to test whether intraspecific split times between refugial populations of butterflies are concentrated in the mid Pleistocene, a patterns that has been found for other herbivorous insect and their parasitoids (Bunnefeld *et al.*, 2018). Nevertheless, our contrasting finding of both gene flow signals in old contact zone pairs (e.g. *Pontia*) and no evidence for gene flow (and complete sympatry) in the youngest pair (*Leptidea*) suggests that the “grey zone of speciation” may be very wide indeed for European butterflies.

### Outlook

Given the challenges of demographic inference from transcriptome data (in particular the high relative recombination rate in butterflies), we have deliberately resisted the temptation to fit explicit models of demographic history. Our goal was instead to establish robust and comparable lower bounds for the age of butterfly sister species in Europe. Being based on mean divergence at 4D sites, these lower bounds for species ages make minimal assumptions and unaffected by recombination. Likewise, we have decided to focus on a simple and conservative diagnostic for introgression.

Delving deeper into the speciation process will require examination of whole-genome data from larger samples under realistic models of speciation history. Fitting explicit models of speciation, ideally including both selection and gene flow, would not only refine estimates for the onset of divergence between recent species but also allow us to quantify the likely end-points (if present) of speciation processes. While it is straightforward to determine lower bounds for the onset of divergence under simple null models that assume no gene flow, as we have done here, estimating upper bounds of species divergence in the presence of gene flow is a much harder inference problem. As pointed out by Barton (Barton & Hewitt, 1985), the initial time of divergence may be unknowable given that post-divergence gene flow eventually erases all information about this parameter. Although current and historic levels of gene flow between European butterfly sister species remain to be determined, our results already suggest that their speciation histories are older and potentially slower than had been assumed by previous phylogeographic studies based on mt data. It will be fascinating to understand the evolutionary forces that drive both this general pattern as well as its exceptions, in particular, the selection responsible for the origin of very young but complete (in terms of reproductive isolation) cryptic species such as *Leptidea* (Talla *et al.*, 2019) and the recently discovered *Spialia rosae* (Hernández-Roldán *et al.*, 2016).

## Acknowledgements

This work was supported by an ERC starting grant (ModelGenomLand). SE is supported by an EastBio studentship from the British Biological Sciences Research Council (BBSRC). KL is supported by a fellowship from the Natural Environment Research Council (NERC, NE/L011522/1). VD is supported by the Academy of Finland (Academy Research Fellow, decision no. 328895). RV acknowledges support from project PID2019-107078GB-I00 (AEI, Spain / ERDF, EU). LD acknowledges support from project “Ricerca e conservazione sugli Impollinatori dell’Arcipelago Toscano e divulgazione sui Lepidotteri del parco”. AH is supported by a Biotechnology and Biological Sciences Research Council (BBSRC) David Phillips Fellowship (BB/N020146/1). We thank Alex Mackintosh and Simon Martin for helpful comments on an earlier draft, Richard Lewington for permission to reproduce his butterfly illustrations Andres de la Filia for help in the molecular lab and Edinburgh Genomics for sequencing.

## Compliance with ethical standards

Field sampling of butterflies was conducted in compliance with the School of Biological Sciences Ethics Committee at the University of Edinburgh and the ERC ethics review procedure. Permissions for field sampling were obtained from the Generalitat de Catalunya (SF/639), the Gobierno de Aragon (INAGA/500201/24/2018/0614 to Karl Wotton) and the Gobierno del Principado de Asturias (014252). The samples for *Z. cassandra* from Elba were collected after permission from the Italian Ministero dell’Ambiente e della Tutela del Territorio e del Mare (Prot. 0012493/ PNM 24/06/2015).

## Author Contributions

SE, DRL, RV, and KL conceived the study; SE performed molecular labwork and analysed the data. DRL, LD, and KL provided tools for analyses. AH, VD, and RV contributed samples. KL, DRL, and MGR supervised the project. SE, KL, and LD wrote the paper with input from all coauthors.

## Data Availability Statement

Sequence alignments and raw read data will be available from a Dryad repository at time of publication. The script used for calculating diversity and divergence is available at https://github.com/samebdon/orthodiver/blob/master/orthodiver.py.

## Supplementary Information

**Figure S1:**
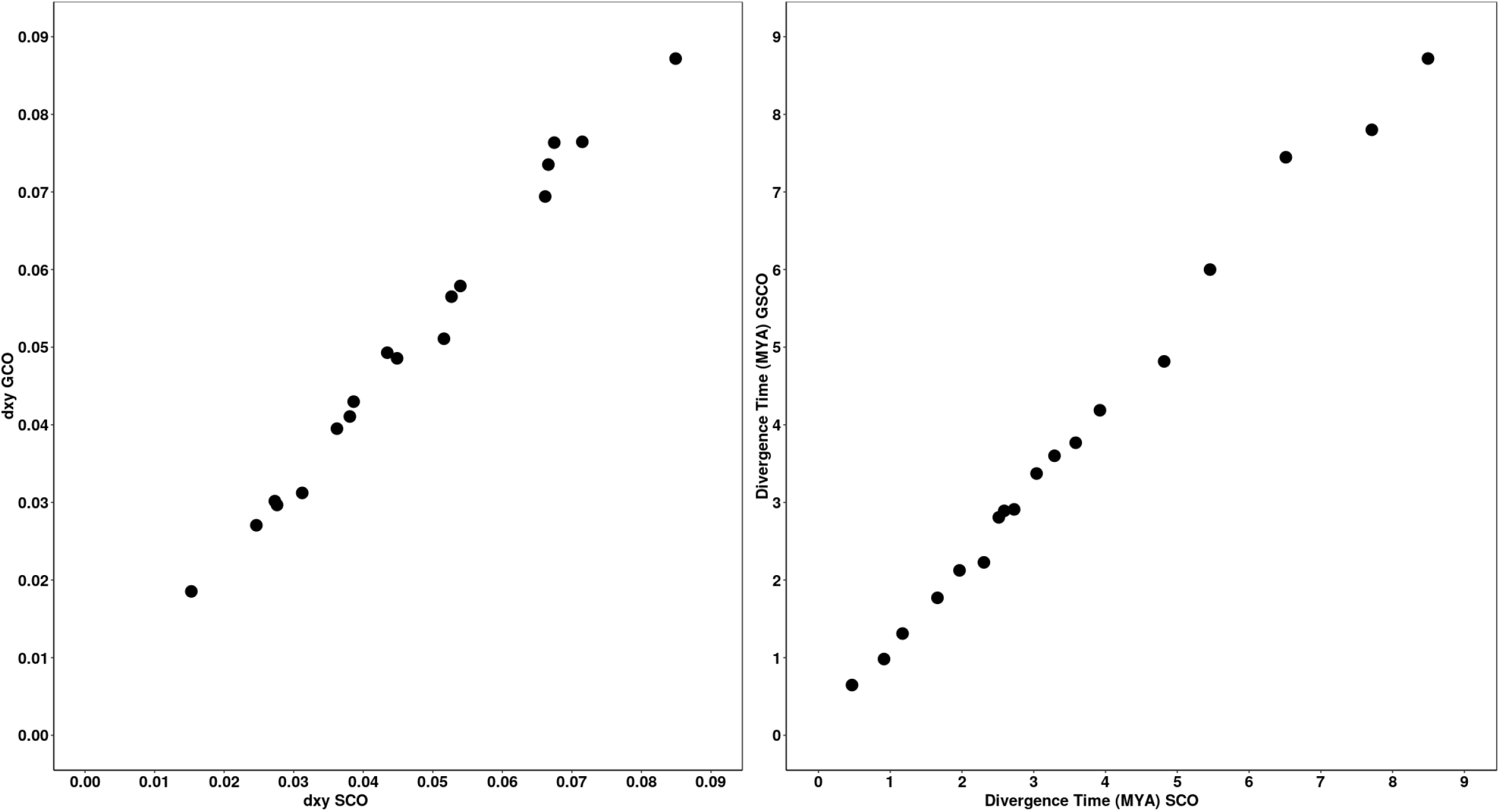
Left) Mean divergence (*d_ry_*) at 4D sites for 18 butterfly species pairs is highly similar at single copy orthologues that are present across all pairs (SCO) and single copy orthologues that are present in each pair/genus (GSCO). Right) Age estimates based on *d_a_* are unaffected by the filtering of orthologues.

**Figure S2:**
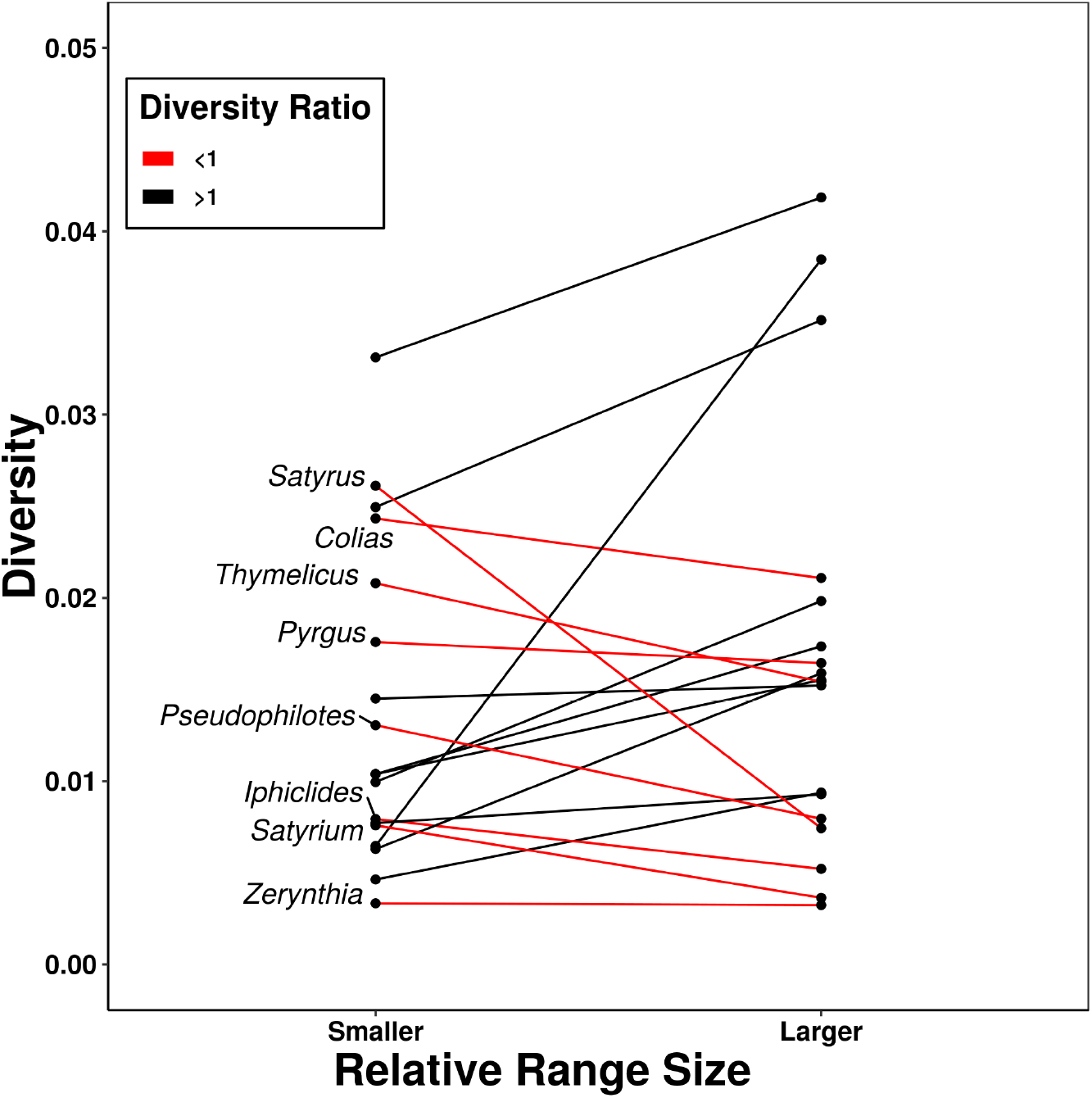
Mean diversity (π) at 4D sites for 18 butterfly species pairs. In most (10) pairs, the species with the smaller range has lower π.

**Figure S3:**
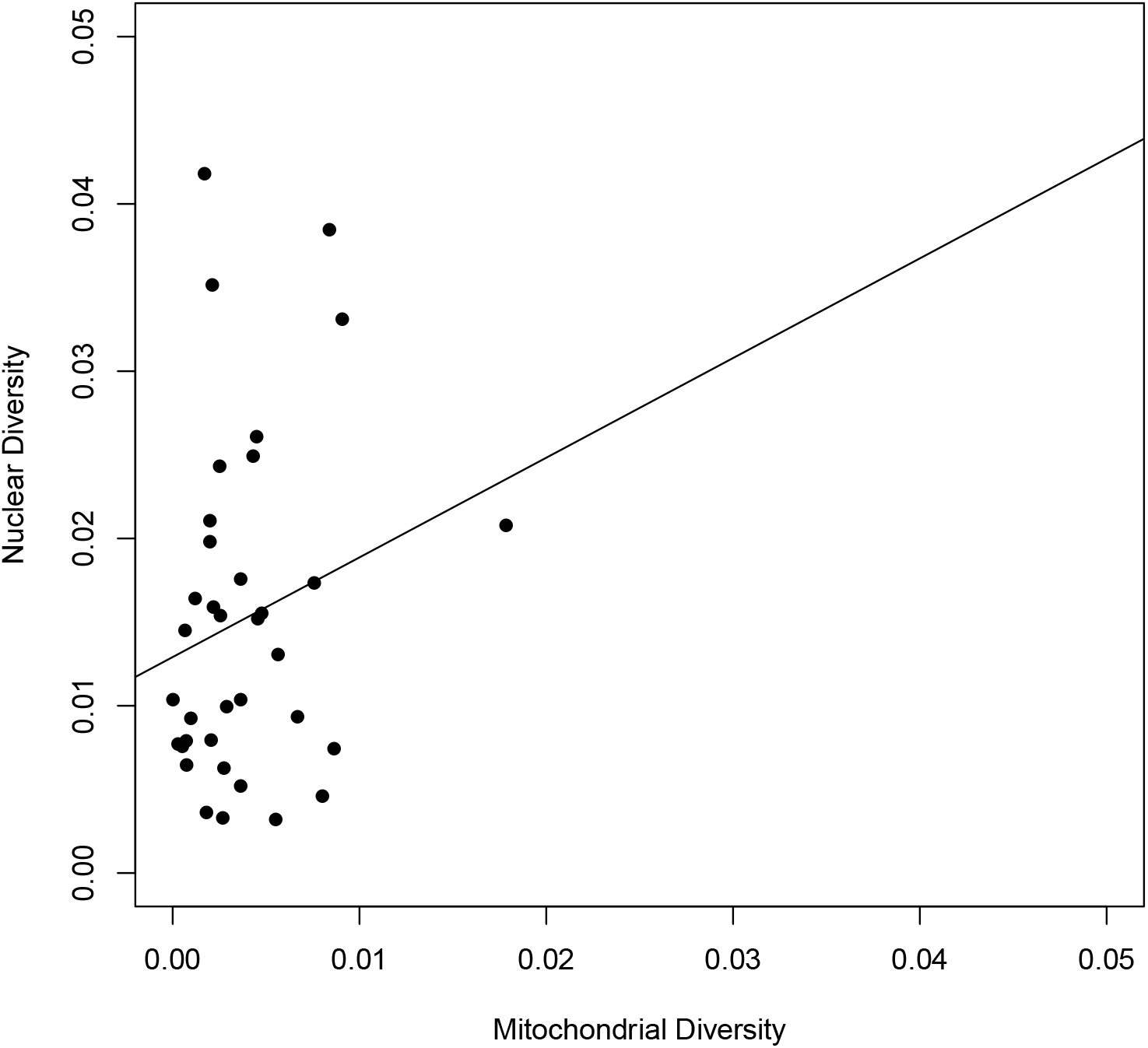
Mitochondrial diversity against nuclear diversity estimated at 4D sites for 36 butterfly species. The slope of best fit is positive (0.07, *R*^2^ = 0.0144) but not significant (t = 1.229, df = 34, p = 0.228).

**Figure S4:**
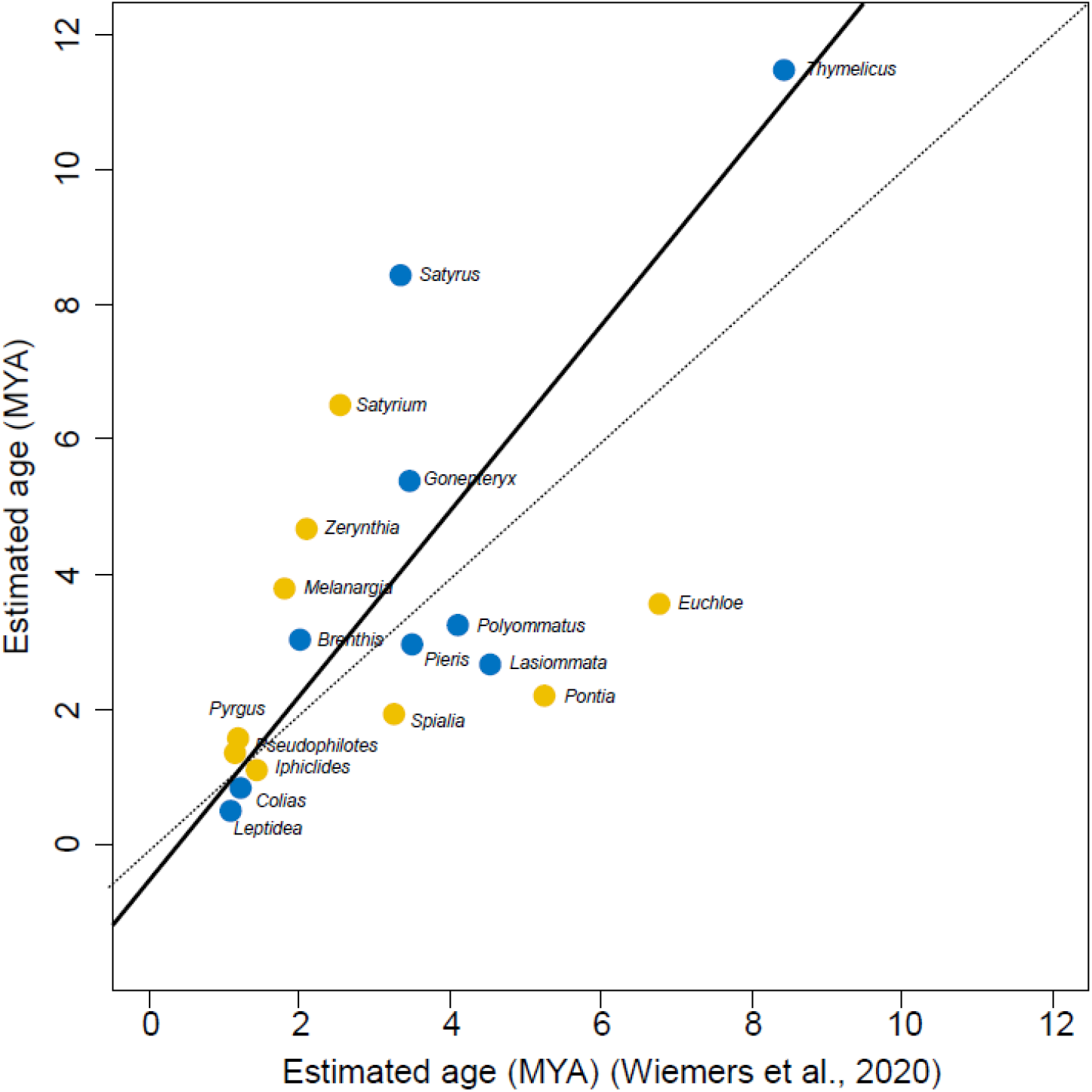
A standardized major axis regression showing a relationship between the age estimates of sister pair nodes in the time calibrated multilocus phylogeny of Wiemers *et al.* (2020) and our estimates from nuclear 4D sites. Yellow data points represent species pairs which abut at contact zones, and blue represents sympatric pairs.

**Figure S5:**
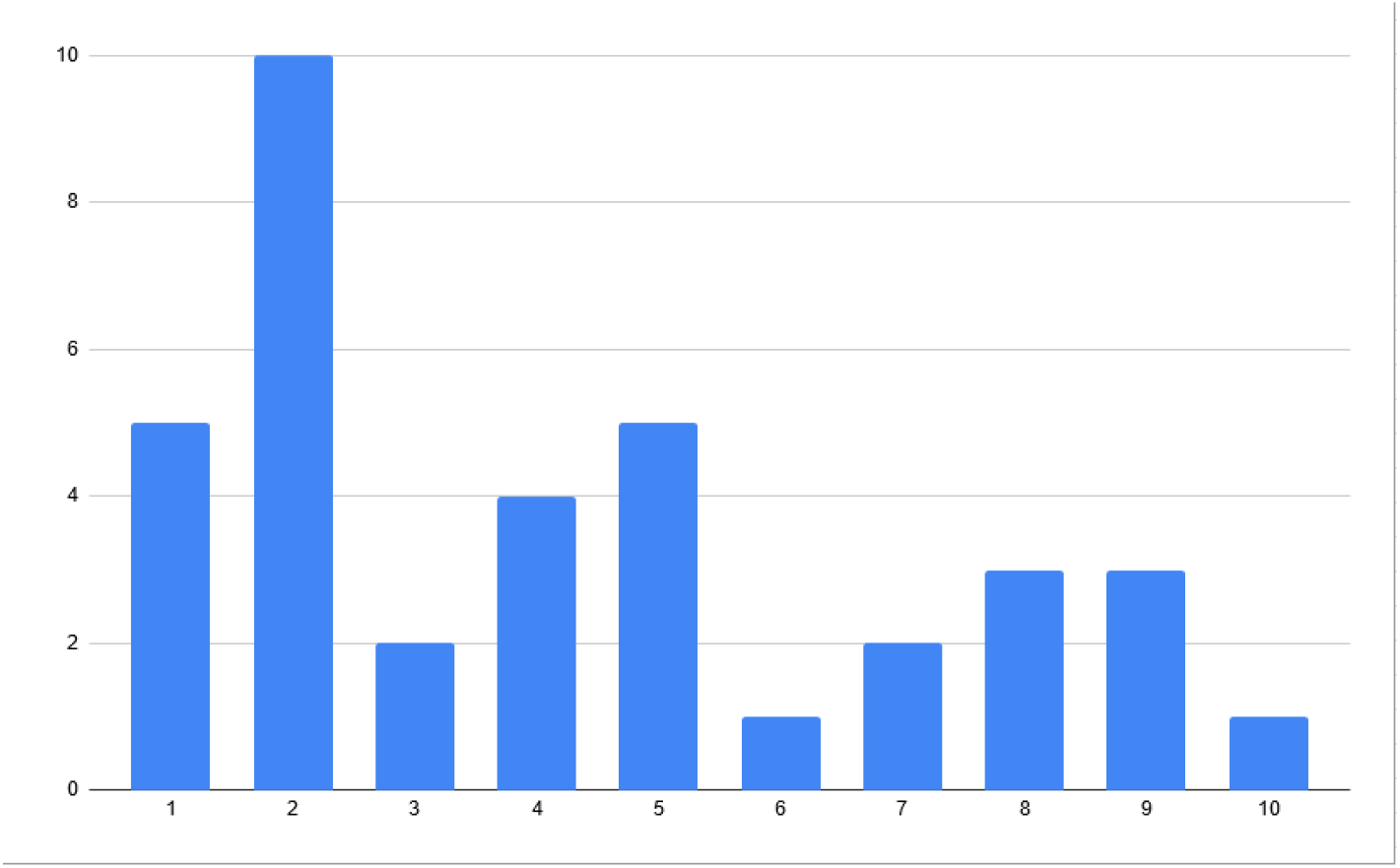
Distribution of the number of loci used by Wiemers *et al.* (2020) for the species used in our study.

**Figure S6:**
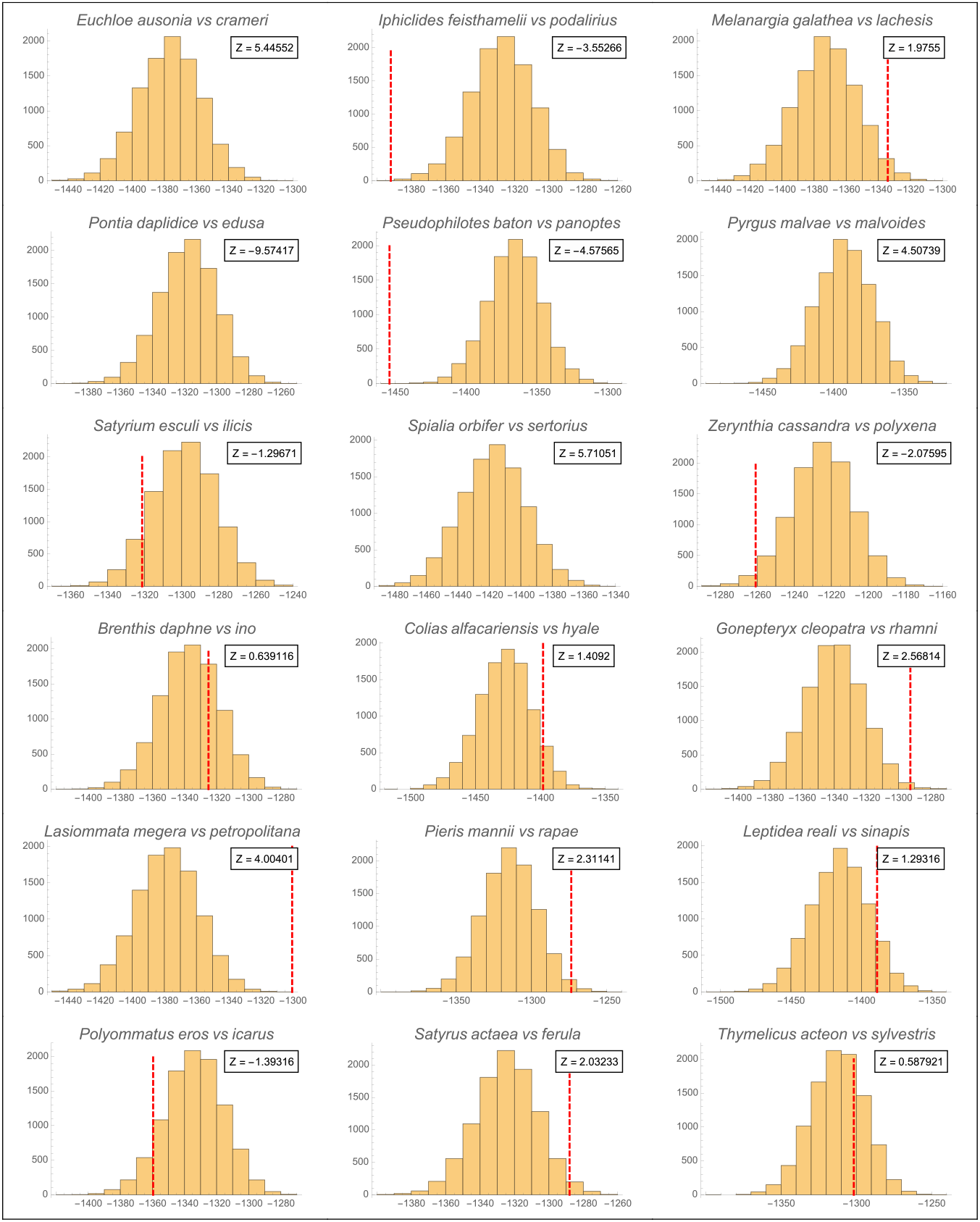
Distribution of log-likelihoods obtained by re-sampling 10,000 datasets from the expected distribution of S for each species pair. The red dashed line is the log-likelihood of the observed data. Data-sets with a Z score greater than 1.96 show narrower S distributions than expected. Data-sets with a Z score less than −1.96 show broader S distributions than expected.

## Notes

### Competing Interest Statement

The authors have declared no competing interest.

### Summary of Updates

Figure 2 revised, contributions and data availability sections appended.

## References

Abascal, F., Zardoya, R. & Telford, M.J. (2010). TranslatorX: multiple alignment of nucleotide sequences guided by amino acid translations. Nucleic Acids Research, 38, W7–W13. ISSN 0305-1048.

Abbott, R.J., Smith, L.C., Milne, R.I., Crawford, R.M., Wolff, K. & Balfour, J. (2000). Molecular analysis of plant migration and refugia in the arctic. Science, 289(5483), 1343–1346.

Andrews, S. et al. (2010). Fastqc: a quality control tool for high throughput sequence data.

Avise, J.C., Walker, D. & Johns, G.C. (1998). Speciation durations and Pleistocene effects on vertebrate phylogeography. Proceedings of the Royal Society of London. Series B: Biological Sciences, 265(1407), 1707–1712.

Bacilieri, R., Ducousso, A., Petit, R.J. & Kremer, A. (1996). Mating system and asymmetric hybridization in a mixed stand of European oaks. Evolution, 50(2), 900–908.

Barraclough, T.G. & Vogler, A.P. (2000). Detecting the geographical pattern of speciation from species-level phylogenies. The American Naturalist, 155(4), 419–434.

Barton, N. (1980). The fitness of hybrids between two chromosomal races of the grasshopper *Podisma pedestris*. Heredity, 45(1), 47–59.

Barton, N.H. & Hewitt, G.M. (1985). Analysis of hybrid zones. Annual review of Ecology and Systematics, 16(1), 113–148.

Bateson, W. (1909). Heredity and variation in modern lights. Cambridge University Press.

Bernatchez, L. & Wilson, C.C. (1998). Comparative phylogeography of nearctic and palearctic fishes. Molecular ecology, 7(4), 431–452.

Bishop, M.P., Björnsson, H., Haeberli, W., Oerlemans, J., Shroder, J.F. & Tranter, M. (2011). Encyclopedia of snow, ice and glaciers. Springer Science & Business Media.

Boursot, P., Din, W., Anand, R., Darviche, D., Dod, B., Von Deimling, F., Talwar, G. & Bonhomme, F. (1996). Origin and radiation of the house mouse: mitochondrial DNA phylogeny. Journal of Evolutionary Biology, 9(4), 391–415.

Bunnefeld, L., Hearn, J., Stone, G.N. & Lohse, K. (2018). Whole-genome data reveal the complex history of a diverse ecological community. Proceedings of the National Academy of Sciences of the United States of America, 115(28), E6507–E6515. ISSN 1091-6490. doi:10.1073/pnas.1800334115.

Butlin, R. & Hewitt, G. (1985). A hybrid zone between *Chorthippus parallelus parallelus* and *Chorthippus parallelus erythropus (Orthoptera*: *Acrididae*): behavioural characters. Biological Journal of the Linnean Society, 26(3), 287–299.

Camacho, C., Coulouris, G., Avagyan, V., Ma, N., Papadopoulos, J., Bealer, K. & Madden, T.L. (2009). Blast+: architecture and applications. BMC bioinformatics, 10(1), 421.

Chen, S., Zhou, Y., Chen, Y. & Gu, J. (2018). fastp: an ultra-fast all-in-one fastq preprocessor. Bioinformatics, 34(17), i884–i890.

Chesser, R.T. & Zink, R.M. (1994). Modes of speciation in birds: a test of Lynch’s method. Evolution, 48(2), 490–497.

Coyne, J.A. & Orr, H.A. (2004). Speciation. sunderland, ma.

Dapporto, L., Cini, A., Vodă, R., Dincă, V., Wiemers, M., Menchetti, M., Magini, G., Talavera, G., Shreeve, T., Bonelli, S., Casacci, L.P., Balletto, E., Scalercio, S. & Vila, R. (2019). Integrating three comprehensive data sets shows that mitochondrial DNA variation is linked to species traits and paleogeographic events in European butterflies. Molecular Ecology Resources, 19(6), 1623–1636. doi:10.1111/1755-0998.13059.

Dapporto, L., Ramazzotti, M., Fattorini, S., Talavera, G., Vila, R. & Dennis, R.L. (2013). recluster: an unbiased clustering procedure for beta-diversity turnover. Ecography, 36(10), 1070–1075.

Dennis, R., Williams, W. & Shreeve, T. (1991). A multivariate approach to the determination of faunal structures among european butterfly species (*Lep-idoptera: Rhopalocera*). Zoological Journal of the Linnean Society, 101(1), 1–49.

Descimon, H. & Mallet, J. (2009). Bad species. Ecology of butterflies in Europe, 500(C), 219.

Dincă, V., Lee, K.M., Vila, R. & Mutanen, M. (2019). The conundrum of species delimitation: a genomic perspective on a mitogenetically super-variable butterfly. Proceedings of the Royal Society B, 286(1911), 20191311.

Dincă, V., Montagud, S., Talavera, G., Hernández-Roldán, J., Munguira, M.L., García-Barros, E., Hebert, P.D.N. & Vila, R. (2015). DNA barcode reference library for Iberian butterflies enables a continental-scale preview of potential cryptic diversity. Scientific reports, 5, 12395. ISSN 2045-2322. doi:10.1038/srep12395.

Dobzhansky, T. (1937). Genetics and the Origin of Species. 11. Columbia university press.

Emms, D.M. & Kelly, S. (2015). Orthofinder: solving fundamental biases in whole genome comparisons dramatically improves orthogroup inference accuracy. Genome biology, 16(1), 157.

Ewels, P., Magnusson, M., Lundin, S. & Käller, M. (2016). Multiqc: summarize analysis results for multiple tools and samples in a single report. Bioinformatics, 32(19), 3047–3048.

Finn, R.D., Clements, J. & Eddy, S.R. (2011). Hmmer web server: interactive sequence similarity searching. Nucleic acids research, 39(suppl_2), W29–W37.

Galtier, N., Nabholz, B., Glémin, S. & Hurst, G. (2009). Mitochondrial DNA as a marker of molecular diversity: a reappraisal. Molecular ecology, 18(22), 4541–4550.

Garrison, E. & Marth, G. (2012). Haplotype-based variant detection from shortread sequencing. arXiv preprint arXiv:1207.3907.

Gaunet, A., Dincă, V., Dapporto, L., Montagud, S., Vodă, R., Schär, S., Badiane, A., Font, E. & Vila, R. (2019a). Two consecutive *Wolbachia*-mediated mitochondrial introgressions obscure taxonomy in palearctic swallowtail butterflies *(Lepidoptera, Papilionidae*). Zoologica Scripta, 48(4), 507–519.

Gaunet, A., Dincă, V.E., Dapporto, L., Montagud, S., Vodă, R., Schör, S., Badiane, A., Font, E. & Vila, R. (2019b). Two consecutive *Wolbachia-mediated* mitochondrial introgressions obscure taxonomy in palearctic swallowtail butterflies *(Lepidoptera, Papilionidae*). Zoologica Scripta. doi:10.1111/zsc.12355.

Gibbard, P. & Head, M.J. (2009). The definition of the Quaternary system/era and the Pleistocene series/epoch. Quaternaire, 20(2), 125–133.

Giordano, R., Jackson, J.J. & Robertson, H.M. (1997). The role of *Wolbachia* bacteria in reproductive incompatibilities and hybrid zones of *Diabrotica* beetles and *Gryllus* crickets. Proceedings of the National Academy of Sciences, 94(21), 11439–11444.

Grabherr, M.G., Haas, B.J., Yassour, M., Levin, J.Z., Thompson, D.A., Amit, I., Adiconis, X., Fan, L., Raychowdhury, R., Zeng, Q. et al. (2011). Full-length transcriptome assembly from rna-seq data without a reference genome. Nature biotechnology, 29(7), 644.

Graham, R.I. & Wilson, K. (2012). Male-killing *Wolbachia* and mitochondrial selective sweep in a migratory African insect. BMC evolutionary biology, 12(1), 204.

Haas, B., Papanicolaou, A. et al. (2016). Transdecoder (find coding regions within transcripts).

Habel, J.C., Meyer, M., El Mousadik, A. & Schmitt, T. (2008). Africa goes europe: The complete phylogeography of the marbled white butterfly species complex *Melanargia galathea/M. lachesis (Lepidoptera*: *Satyridae*). Organisms Diversity & Evolution, 8(2), 121–129.

Habel, J.C., Vila, R., Vodaa, R., Husemann, M., Schmitt, T. & Dapporto, L. (2017). Differentiation in the marbled white butterfly species complex driven by multiple evolutionary forces. Journal of Biogeography, 44(2), 433–445.

Haffer, J. (1969). Speciation in amazonian forest birds. Science, 165(3889), 131–137.

Hajibabaei, M., Janzen, D.H., Burns, J.M., Hallwachs, W. & Hebert, P.D. (2006). DNA barcodes distinguish species of tropical *Lepidoptera*. Proceedings of the National Academy of Sciences, 103(4), 968–971.

Hansen, J., Sato, M., Russell, G. & Kharecha, P. (2013). Climate sensitivity, sea level and atmospheric carbon dioxide. Philosophical Transactions of the Royal Society A: Mathematical, Physical and Engineering Sciences, 371(2001), 20120294.

Hernández-Roldán, J.L., Dapporto, L., Dincă, V., Vicente, J.C., Hornett, E.A., Síchová, J., Lukhtanov, V.A., Talavera, G. & Vila, R. (2016). Integrative analyses unveil speciation linked to host plant shift in *Spialia* butterflies. Molecular Ecology, 25(17), 4267–4284.

Hewitt, G. (2000). The genetic legacy of the Quaternary ice ages. Nature, 405(6789), 907–913.

Hewitt, G.M. (1996). Some genetic consequences of ice ages, and their role in divergence and speciation. Biological journal of the Linnean Society, 58(3), 247–276.

Hewitt, G.M. (1999). Post-glacial re-colonization of European biota. Biological journal of the Linnean Society, 68(1-2), 87–112.

Hewitt, G.M. (2001). Speciation, hybrid zones and phylogeography—or seeing genes in space and time. Molecular ecology, 10(3), 537–549.

Hewitt, G.M. (2011). Quaternary phylogeography: the roots of hybrid zones. Genetica, 139(5), 617–638.

Hinojosa, J.C., Koubínová, D., Szenteczki, M.A., Pitteloud, C., Dincă, V., Alvarez, N. & Vila, R. (2019). A mirage of cryptic species: Genomics uncover striking mitonuclear discordance in the butterfly *Thymelicus sylvestris*. Molecular ecology, 28(17), 3857–3868.

Hofreiter, M. & Stewart, J. (????). Ecological change, range fluctuations and population dynamics during the Pleistocene. Current Biology, (14), R584–R594. ISSN 0960-9822. doi:10.1016/J.CUB.2009.06.030.

Hurst, G.D. & Jiggins, F.M. (2005). Problems with mitochondrial DNA as a marker in population, phylogeographic and phylogenetic studies: the effects of inherited symbionts. Proceedings of the Royal Society B: Biological Sciences, 272(1572), 1525–1534.

Jiggins, F.M. (2003). Male-killing *Wolbachia* and mitochondrial DNA: Selective sweeps, hybrid introgression and parasite population dynamics. Genetics, 164(1), 5–12.

Jombart, T. & Dray, S. (2010). adephylo: exploratory analyses for the phylogenetic comparative method. Bioinformatics, 26, 1907–1909. doi:10.1093/bioinformatics/btq292.

Kaliontzopoulou, A., Pinho, C., Harris, D.J. & Carretero, M.A. (2011). When cryptic diversity blurs the picture: a cautionary tale from Iberian and north African *Podarcis* wall lizards. Biological Journal of the Linnean Society, 103(4), 779–800.

Keightley, P.D., Ness, R.W., Halligan, D.L. & Haddrill, P.R. (2014a). Estimation of the spontaneous mutation rate per nucleotide site in a *Drosophila melanogaster* full-sib family. Genetics, 196(1), 313–320. ISSN 0016-6731. doi:10.1534/genetics.113.158758.

Keightley, P.D., Pinharanda, A., Ness, R.W., Simpson, F., Dasmahapatra, K.K., Mallet, J., Davey, J.W. & Jiggins, C.D. (2014b). Estimation of the spontaneous mutation rate in heliconius melpomene. Molecular biology and evolution, 32(1), 239–243.

Klicka, J. & Zink, R.M. (1997). The importance of recent ice ages in speciation: A failed paradigm. Science, 277(5332), 1666–1669. ISSN 0036-8075. doi:10.1126/science.277.5332.1666.

Knowles, L.L. & Carstens, B.C. (2007). Delimiting species without monophyletic gene trees. Systematic biology, 56(6), 887–895.

Kodandaramaiah, U., Simonsen, T.J., Bromilow, S., Wahlberg, N. & Sperling, F. (2013). Deceptive single-locus taxonomy and phylogeography: *Wolbachia*-associated divergence in mitochondrial DNA is not reflected in morphology and nuclear markers in a butterfly species. Ecology and Evolution, 3(16), 5167–5176.

Krijgsman, W., Hilgen, F., Raffi, I., Sierro, F. & Wilson, D. (1999). Chronology, causes and progression of the Messinian salinity crisis. Nature, 400(6745), 652–655.

Kruuk, L.E., Gilchrist, J.S. & Barton, N.H. (1999). Hybrid dysfunction in fire-bellied toads *(Bombina*). Evolution, 53(5), 1611–1616.

Kudrna, O. (2019). Distribution of butterflies and skippers in Europe: (Lepidoptera: Rhopalocera, Grypocera): 24 years mapping European butterflies (1995-2019): final report. Spolecnost pro Ochranu Motylu (SOM).

Kudrna, O., Harpke, A., Lux, K., Pennerstorfer, J., Schweiger, O., Settele, J. & Wiemers, M. (2011). Distribution atlas of butterflies in Europe. Gesellschaft fur Schmetterlingsschutz.

Lai, B.C.G. & Pullin, A.S. (2004). Phylogeography, genetic diversity and conservation of the large copper butterfly *Lycaena dispar* in Europe. Journal of Insect Conservation, 8(1), 27–36.

Li, H. (2011). A statistical framework for snp calling, mutation discovery, association mapping and population genetical parameter estimation from sequencing data. Bioinformatics, 27(21), 2987–2993.

Li, H. (2013). Aligning sequence reads, clone sequences and assembly contigs with bwa-mem. arXiv preprint arXiv:1303.3997.

Librado, P. & Rozas, J. (2009). DnaSP v5: a software for comprehensive analysis of DNA polymorphism data. Bioinformatics, 25(11), 1451–1452.

Lohse, K., Harrison, R.J. & Barton, N.H. (2011). A general method for calculating likelihoods under the coalescent process. Genetics, 189(3), 977–987. ISSN 0016-6731. doi:10.1534/genetics.111.129569.

Lorkovic, Z. (1973). 150 jahre bis zur entdeckung der präimaginalstadien von *Spialia orbifer* hbn.(Lep., *Hesperiidae*). Acta Entomologica Yugoslavica, 9, 67–70.

Mackintosh, A., Laetsch, D.R., Hayward, A., Charlesworth, B., Waterfall, M., Vila, R. & Lohse, K. (2019). The determinants of genetic diversity in butterflies. Nature Communications, 10(1), 3466. ISSN 2041-1723. doi:10.1038/s41467-019-11308-4.

Martin, S.H., Davey, J.W., Salazar, C. & Jiggins, C.D. (2019). Recombination rate variation shapes barriers to introgression across butterfly genomes. PLoS biology, 17(2), e2006288.

Martin, S.H., Möst, M., Palmer, W.J., Salazar, C., McMillan, W.O., Jiggins, F.M. & Jiggins, C.D. (2016). Natural selection and genetic diversity in the butterfly *Heliconius melpomene*. Genetics, 203(1), 525–541.

Martin, S.H., Singh, K.S., Gordon, I.J., Omufwoko, K.S., Collins, S., Warren, I.A., Munby, H., Brattström, O., Traut, W., Martins, D.J. et al. (2020). Whole-chromosome hitchhiking driven by a male-killing endosymbiont. PLoS biology, 18(2), e3000610.

Mayr, E. (1947). Ecological factors in speciation. Evolution, 1(4), 263–288.

McKenna, A., Hanna, M., Banks, E., Sivachenko, A., Cibulskis, K., Kernytsky, A., Garimella, K., Altshuler, D., Gabriel, S., Daly, M. et al. (2010). The genome analysis toolkit: a mapreduce framework for analyzing next-generation dna sequencing data. Genome research, 20(9), 1297–1303.

Muller, H. (1942). Isolating mechanisms, evolution, and temperature. In Biol. Symp., volume 6, pages 71–125.

Nei, M. & Li, W.H. (1979). Mathematical model for studying genetic variation in terms of restriction endonucleases. Proceedings of the National Academy of Sciences, 76(10), 5269–5273. ISSN 0027-8424. doi:10.1073/pnas.76.10.5269.

Nolen, Z., B, Y., Irisarri, I., Liu S, Groot, C., Amby, D., Mayer, F., Gilbert, M. & R, P. (2020). Historical isolation facilitates species radiation by sexual selection: insights from *Chorthippus* grasshoppers. Authorea.

Nürnberger, B., Lohse, K., Fijarczyk, A., Szymura, J.M. & Blaxter, M.L. (2016). Para-allopatry in hybridizing fire-bellied toads (*Bombina bombina* and *B. var-iegata*): Inference from transcriptome-wide coalescence analyses. Evolution, 70(8), 1803–1818.

Pigot, A.L. & Tobias, J.A. (2013). Species interactions constrain geographic range expansion over evolutionary time. Ecology letters, 16(3), 330–338.

Platania, L., Vodă, R., Dincă, V., Talavera, G., Vila, R. & Dapporto, L. (2020). Integrative analyses on western palearctic *Lasiommata* reveal a mosaic of nascent butterfly species. Journal of Zoological Systematics and Evolutionary Research.

Porter, A.H., Wenger, R., Geiger, H., Scholl, A. & Shapiro, A.M. (1997). The *Pontia daplidice-edusa* hybrid zone in northwestern Italy. Evolution, 51(5), 1561–1573.

Quinlan, A.R. & Hall, I.M. (2010). Bedtools: a flexible suite of utilities for comparing genomic features. Bioinformatics, 26(6), 841–842.

Ratnasingham, S. & Hebert, P.D. (2007). Bold: The barcode of life data system (http://www.barcodinglife.org). Molecular ecology notes, 7(3), 355–364.

Ritter, S., Michalski, S.G., Settele, J., Wiemers, M., Fric, Z.F., Sielezniew, M., Saŝiá, M., Rozier, Y. & Durka, W. (2013). *Wolbachia* infections mimic cryptic speciation in two parasitic butterfly species, *Phengaris teleius* and *P. nausithous (Lepidoptera*: *Lycaenidae*). PLoS One, 8(11), e78107.

Rokas, I. (2000). *Wolbachia* as a speciation agent. Trends in Ecology & Evolution, 15(2), 44.

Roux, C., Fraisse, C., Romiguier, J., Anciaux, Y., Galtier, N. & Bierne, N. (2016). Shedding light on the grey zone of speciation along a continuum of genomic divergence. PLoS biology, 14(12), e2000234.

Schmitt, T. (2007a). Molecular biogeography of europe: Pleistocene cycles and postglacial trends. Frontiers in zoology, 4(1), 11.

Schmitt, T. (2007b). Molecular biogeography of Europe: Pleistocene cycles and postglacial trends. Frontiers in zoology, 4(1), 11.

Schoville, S.D., Roderick, G.K. & Kavanaugh, D.H. (2012). Testing the ‘Pleistocene species pump’in alpine habitats: lineage diversification of flightless ground beetles *(Coleoptera*: *Carabidae*: *Nebria*) in relation to altitudinal zonation. Biological Journal of the Linnean Society, 107(1), 95–111.

Shoemaker, D.D., Katju, V. & Jaenike, J. (1999). *Wolbachia* and the evolution of reproductive isolation between *Drosophila recens* and *Drosophila subquinaria*. Evolution, 53(4), 1157–1164.

Simao, F.A., Waterhouse, R.M., Ioannidis, P., Kriventseva, E.V. & Zdobnov, E.M. (2015). Busco: assessing genome assembly and annotation completeness with single-copy orthologs. Bioinformatics, 31(19), 3210–3212.

Skibinski, D. & Beardmore, J. (1979). A genetic study of intergradation between *Mytilus edulis* and *Mytilus galloprovincialis*. Experientia, 35(11), 1442–1444.

Spooner, L. & Ritchie, M. (2006). An unusual phylogeography in the bushcricket *Ephippiger ephippiger* from southern France. Heredity, 97(6), 398–408.

Talla, V., Johansson, A., Dincă, V., Vila, R., Friberg, M., Wiklund, C. & Backström, N. (2019). Lack of gene flow: narrow and dispersed differentiation islands in a triplet of *Leptidea* butterfly species. Molecular ecology, 28(16), 3756–3770.

Talla, V., Suh, A., Kalsoom, F., Dincă, V., Vila, R., Friberg, M., Wiklund, C. & Backström, N. (2017). Rapid increase in genome size as a consequence of transposable element hyperactivity in wood-white *(Leptidea*) butterflies. Genome Biology and Evolution, 9(10), 2491–2505.

Thomas, J.A., Welch, J.J., Lanfear, R. & Bromham, L. (2010). A generation time effect on the rate of molecular evolution in invertebrates. Molecular Biology and Evolution, 27(5), 1173–1180. ISSN 0737-4038.

Todisco, V., Gratton, P., Cesaroni, D. & Sbordoni, V. (2010). Phylogeography of *Parnassius apollo*: hints on taxonomy and conservation of a vulnerable glacial butterfly invader. Biological Journal of the Linnean Society, 101(1), 169–183.

Toews, D.P. & Brelsford, A. (2012). The biogeography of mitochondrial and nuclear discordance in animals. Molecular Ecology, 21(16), 3907–3930.

Tolman, T. & Lewington, R. (2013). Collins British butterfly guide. Collins.

Vodă, R., Dapporto, L., Dincă, V. & Vila, R. (2015). Why do cryptic species tend not to co-occur? a case study on two cryptic pairs of butterflies. PloS one, 10(2), e0117802.

Wall, J.D. (2003). Estimating ancestral population sizes and divergence times. Genetics, 163(1), 395–404. ISSN 0016-6731.

Wang, Y. & Hey, J. (2010). Estimating divergence parameters with small samples from a large number of loci. Genetics, 184(2), 363–379.

Weir, J.T. & Price, T.D. (2011). Limits to speciation inferred from times to secondary sympatry and ages of hybridizing species along a latitudinal gradient. The American Naturalist, 177(4), 462–469.

Wiemers, M., Balletto, E., Dincă, V., Fric, Z.F., Lamas, G., Lukhtanov, V., Munguira, M.L., van Swaay, C., Vila, R., Vliegenthart, A., Wahlberg, N. & Verovnik, R. (2018). An updated checklist of the european butterflies *(Lepidoptera, Papilionoidea*). ZooKeys, 811, 9–45. ISSN 1313-2989.

Wiemers, M., Chazot, N., Wheat, C.W., Schweiger, O. & Wahlberg, N. (2020). A complete time-calibrated multi-gene phylogeny of the European butterflies. ZooKeys, 938, 97.

Wiemers, M., Stradomsky, B.V. & Vodolazhsky, D.I. (2010). A molecular phylogeny of *Polyommatus s. str.* and *Plebicula* based on mitochondrial COI and nuclear ITS2 sequences *(Lepidoptera*: *Lycaenidae*). European Journal of Entomology, 107(3), 325.

Wilkinson-Herbots, H.M. (2012). The distribution of the coalescence time and the number of pairwise nucleotide differences in a model of population divergence or speciation with an initial period of gene flow. Theoretical population biology, 82(2), 92–108.

Wohlfahrt, T. (1996). Vergleichende untersuchun gen über die größe und form der augenflecken am analende der hinterflügel von *Iphiclides podalirius podalirius* (linnaeus, 1758) und I. podalirius feisthamelii (duponchel, 1832). 19, 281–288.

Zinetti, F., Dapporto, L., Vovlas, A., Chelazzi, G., Bonelli, S., Balletto, E. & Ciofi, C. (2013). When the rule becomes the exception. no evidence of gene flow between two *Zerynthia* cryptic butterflies suggests the emergence of a new model group. PLoS One, 8(6), e65746.

